# Drosophila Rif1 is critical for repair following P-element excision and influences pathway choice at double-strand breaks

**DOI:** 10.1101/2022.05.24.493293

**Authors:** Justin R. Blanch, Manan Krishnamurthy, Jacob T. Zuckerman, Mitch McVey

**Author notes:** these authors contributed equally to this work (Justin R. Blanch, Manan Krishnamurthy).

## Abstract

Rif1 plays important roles in the repair of DNA double-strand breaks in multiple organisms. In mammals, RIF1 promotes non-homologous end joining and suppresses homologous recombination by interacting with 53BP1 to inhibit resection. In *Saccharomyces cerevisiae*, Rif1 directly binds DNA to inhibit resection and promote non-homologous end-joining. Yeast Rif1 can also facilitate long-range resection and promote single-strand annealing. Since it is not clear if Rif1 regulates resection-mediated pathway choice in other eukaryotes, we explored the role of Rif1 in double-strand break repair in *Drosophila melanogaster*. We found that *rif1* mutants are not sensitive to ionizing radiation or hydroxyurea, demonstrating that it is not essential for the resolution of DNA damage in Drosophila. However, we show that *rif1* null mutants are largely unable to repair a specific type of double-strand break that is induced upon the excision of a P-element transposon. Furthermore, assessment of repair pathway choice at I-*Sce*I-induced breaks revealed Rif1 suppresses homologous recombination and promotes single-strand annealing. Collectively, our findings illustrate Drosophila Rif1 shares functions with both its yeast and mammalian counterparts and serves a unique role in repairing P-element-induced double-strand breaks.

## Introduction

When disruptive DNA double-strand breaks (DSBs) arise in the genome, the selection of a repair pathway greatly influences the genomic outcome. Accurate repair requires specific suites of cellular machinery within distinct phases of the cell cycle, while inaccurate or unsuccessful repair can lead to mutations or cell death. Since multiple pathways can resolve the same DSB and each type varies in accuracy, the selection and timing of repair pathways are highly regulated processes.

Repair pathway choice greatly depends on the extent of 5’ to 3’ DNA resection at the DSB. Extensive resection normally initiates homologous recombination (HR), which involves strand invasion into a homologous template, DNA synthesis, and ligation to complete repair (Jasin and Rothstein 2013; Fugger and West 2016) (Figure 1). HR serves as an accurate method to resolve a break because a sister chromatid is often used as the template for repair. Conversely, inhibition of resection results in religation of DSB ends by non-homologous end-joining (NHEJ), which can also be error-free or causes small insertions or deletions (Weterings and Chen 2008; Wyatt *et al*. 2016; Bhargava *et al*. 2018; Cejka and Symington 2021). Together, HR and NHEJ account for the majority of repair in the cell cycle: NHEJ is favored during G1, while HR is favored during S and G2 (Ceccaldi *et al*. 2016; Arnoult *et al*. 2017; Chang *et al*. 2017; Zhao *et al*. 2017). Despite these preferences, NHEJ and HR genes are often co-expressed in G1/S (Mjelle *et al*. 2015). Therefore, exploring proteins that control resection-mediated choice is critical for understanding the regulation of both pathways.

**Figure 1.**
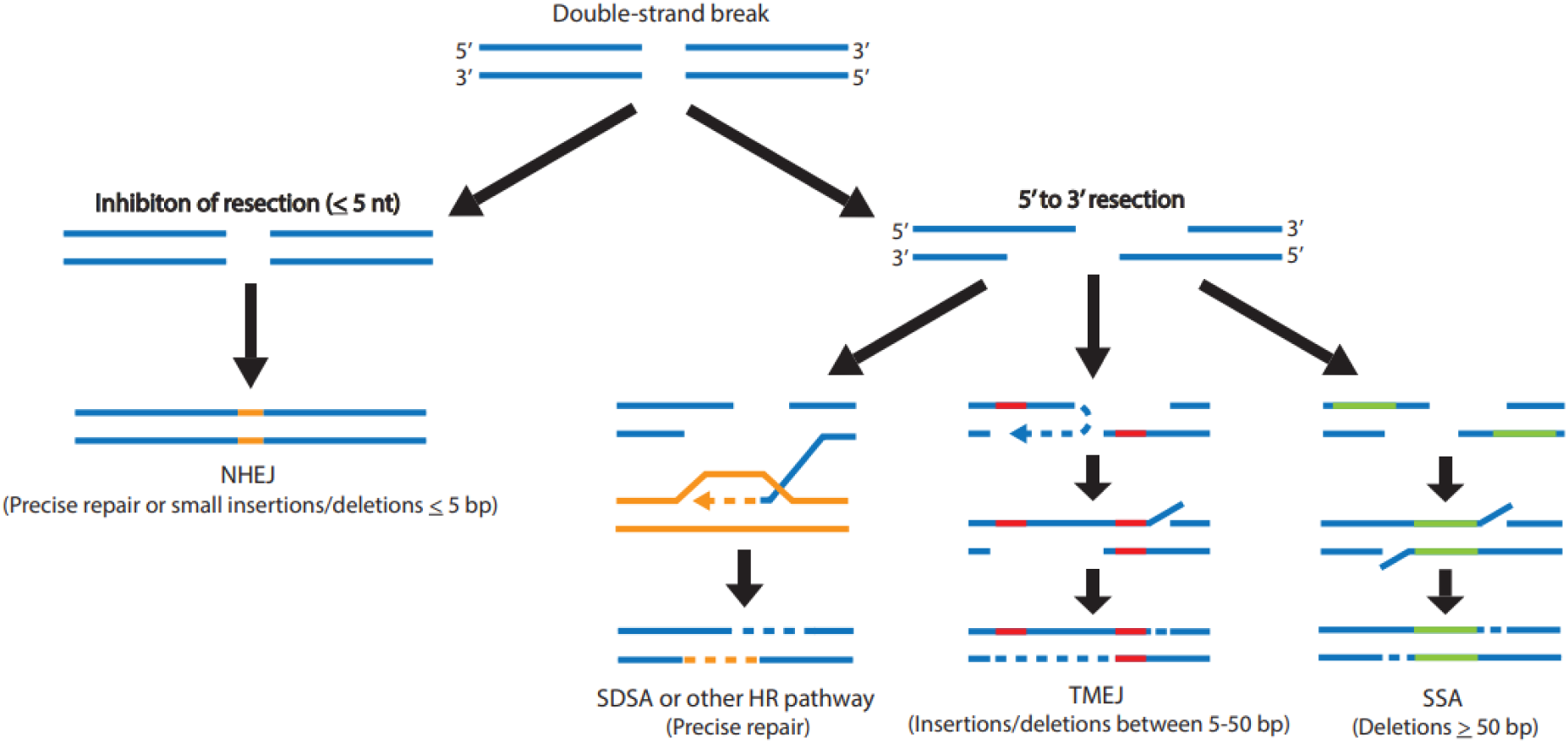
Double-strand break repair pathways. Non-homologous end-joining (NHEJ) occurs prior to resection of the break, while homologous recombination (HR) is stimulated by resection. Synthesis-dependent strand annealing (SDSA) promotes accurate repair using a sister chromatid or other homologous repair template. Theta-mediated end joining (TMEJ) is a form of alternative end-joining that involves the annealing of short microhomologies and creates small insertions and deletions near the break site. Single-strand annealing (SSA) involves extensive resection, annealing of large homologous sequences, and creates large deletions.

Less prevalent, but more error-prone pathways are also used to repair DSBs. These pathways are often upregulated in cancers and are also governed by resection at the DSB (Blasiak 2021; Ramsden *et al*. 2022). Error-prone theta-mediated end-joining (TMEJ) involves limited resection, annealing of microhomologies, and the generation of small deletions or insertions that are typically larger than mutations found in NHEJ (Figure 1) (Khodaverdian *et al*. 2017; Schimmel *et al*. 2019; Carvajal-Garcia *et al*. 2020; Hanscom and Mcvey 2020; Ramsden *et al*. 2022). Another error-prone pathway, single-strand annealing (SSA), involves extensive resection, annealing of longer homologous regions than TMEJ, and can result in very large deletions ≥ 50 base pairs (bp) (Bhargava *et al*. 2016; Kelso *et al*. 2019). Repair by TMEJ or SSA is often responsible for mutations in precancerous cells deficient in NHEJ or HR resection machinery (Ahrabi *et al*. 2016; Bakr *et al*. 2016). This elevated potential for mutagenicity and malignant transformation underscores the importance of proper regulation of NHEJ and HR resection machinery.

In mammals, resection is largely controlled by the interaction between 53BP1 and RIF1 at the break-site (Chapman *et al*. 2013; Di Virgilio *et al*. 2013; Escribano-Díaz *et al*. 2013; Zimmermann *et al*. 2013). RIF1 complexes with 53BP1 in G1 to prevent end-resection by the BRCA1-CtIP complex and promotes repair by NHEJ (Chapman *et al*. 2013; Escribano-Díaz *et al*. 2013; Bakr *et al*. 2016; Setiaputra *et al*. 2022). The efficacy of RIF1-53BP1 in inhibiting resection depends on two downstream effectors, PP1 and shieldin. PP1 forms a complex with RIF1 to prevent MRN resection machinery from creating single-stranded DNA (ssDNA) (Isobe *et al*. 2021), and shieldin prevents the formation of RPA nucleofilaments (Gupta *et al*. 2018; Setiaputra and Durocher 2019). By means of these various functional modes, RIF1-53BP1 represses resection and governs DSB repair pathway choice between HR and NHEJ in mammals.

Several domains within mammalian RIF1 contribute to its functions in DSB repair and are relatively well-conserved across eukaryotes (Sreesankar *et al*. 2012; Fontana *et al*. 2018). A HEAT repeat domain within the N-terminus of mammalian RIF1 promotes localization of RIF1 to DSBs (Escribano-Díaz *et al*. 2013). A DNA binding domain within the C-terminus helps recruit RIF1 to stalled replication forks and interacts with BLM helicase (Xu *et al*. 2010). BLM normally resolves Holliday Junctions in HR and secondary DNA structures at stalled replication forks (Kaur *et al*. 2021). Depletion of RIF1 disrupts BLM localization to DSBs in mammals, suggesting RIF1 has additional roles in signaling for several types of repair at breaks (Feng *et al*. 2013). Further, a SILK-RVxF motif within the C-terminus interacts with PP1 in Drosophila and the PP1-binding motif inhibits end-resection in mammals (Sreesankar *et al*. 2015; Isobe *et al*. 2021). The PP1-binding motif may be critical for mammalian RIF1 to inhibit resection in NHEJ.

In contrast to its function in mammals, Rif1 can both inhibit and activate resection in budding yeast. One study reported that Rif1 inhibits resection 0.7-4.2 kilobases (kb) away from the break to promote NHEJ in budding yeast (Mattarocci *et al*. 2017). A hook domain within the N-terminus of Rif1 is critical for this inhibition (Mattarocci *et al*. 2017). A separate study reported the loss of budding yeast Rif1 exacerbates DSB resection in *exo1* and *sae2* mutants and reduces levels of resection greater than 2 kb (Martina *et al*. 2014). Further, Rif1 promotes SSA 25 kb away from the break and counteracts the binding of the Rad9 resection inhibitor (Martina *et al*. 2014).

One major difference between the budding yeast and mammalian Rif1 pathways is the emergence of the RIF1-53BP1 interaction in mammals. Yeast do not have a structural homolog of 53BP1 and rather than complexing with another protein, Rif1 directly binds to chromatin around DSB ends (Martina *et al*. 2014). It is unclear if the RIF1-53BP1 interaction is responsible for the difference between mammalian and yeast Rif1-dependent control of resection, and further experiments are necessary to investigate the role of Rif1-mediated resection in other eukaryotes.

Given the varied and somewhat contrasting roles of Rif1 in the regulation of resection in mammals and yeast, we wanted to explore the function of Rif1 in Drosophila. Drosophila Rif1 domains are not entirely conserved with either mammalian or yeast Rif1 domains (Sreesankar *et al*. 2012; Mattarocci *et al*. 2016) and a 53BP1 ortholog has not been identified in Drosophila. Recent studies show the absence of Drosophila Rif1 does not affect viability, but it does lead to modest or substantial reductions in egg hatching frequencies (Munden *et al*. 2018; Seller and O’Farrell 2018). Although Drosophila *Rif1* is expressed throughout embryonic development, constitutive overexpression of *Rif1* causes pupal lethality and abnormal condensation of chromatin (Sreesankar *et al*. 2015). Additionally, Drosophila Rif1 has several important functions in DNA replication. Rif1 extends S-phase during embryonic development (Seller and O’Farrell 2018) and localizes to replication forks to regulate fork progression, copy number, and replication timing (Munden *et al*. 2018; Armstrong *et al*. 2020; Kolesnikova *et al*. 2020). Interestingly, Drosophila Rif1 does not colocalize with γ-H2AvD foci upon inhibition of the cell cycle or induction of DNA damage in cells (Sreesankar *et al*. 2012), which suggests Rif1 may not be directly recruited to DSBs in Drosophila. However, whether Drosophila Rif1 has a role in repairing DNA damage or repair pathway choice has not been reported at the organismal level.

To test the importance of Rif1 in DNA repair in Drosophila, we conducted mutagen sensitivity assays, monitored egg development and hatching, and performed site-specific double-strand break repair assays. We find that Rif1 is not required to repair exogenously-induced damage or breaks that arise in follicle cells prior to egg hatching. In contrast, it is critical for the repair of P-element-induced DSBs. Additionally, Rif1 suppresses HR and promotes SSA at I-*Sce*I-induced breaks. Overall, our data suggest that Drosophila Rif1 participates in mechanisms that govern repair pathway choice, similar to its roles in other organisms.

## Materials and Methods

### Mutant fly stocks

Flies were kept at 25°C and given a standard cornmeal diet. The *rif1*^*null*^ stocks (*rif1*^*1*^ and *rif1*^*2*^) used in this study were gifts from Jared Nordman and were generated by CRISPR-induced mutagenesis (Munden *et al*. 2018). *rif1*^*1*^ contains a 3,864 bp deletion and *rif1*^*2*^ contains a 2,053 bp deletion; both mutations are in the *Rif1* coding sequence and eliminate protein translation.

The *rif1*^*1-7*^ *null* mutant was recovered from an imprecise excision of the *P{EP}Rif1*^*G18022*^ element (Bloomington stock #27427) which produced a large indel and premature stop codons in the *Rif1* coding sequence. A *Δ2-3* transposase source on the second chromosome was used to generate the excision (*CyO, H{w+, Δ2–3*}, from Bloomington stock #2078). The indel was validated by PCR and sequencing (*Rif1* -402 forward, 5’ ATCCAATTGAGTCCCGCCAG 3’; *Rif1* 1958 reverse, 5’ GGTTGCGGAGGGTGTTT AAAC 3’).

### Mutagen sensitivity assays

In hydroxyurea assays, five virgin females homozygous for one *rif1*^*null*^ allele (*rif1*^*1*^*/rif1*^*1*^ or *rif1*^*2*^*/rif1*^*2*^) were crossed with three males heterozygous for the same *rif1*^*null*^ allele (*rif1*^*1*^*/CyO* or *rif1*^*2*^*/CyO*) in each vial for three days at 25°C. On the third day, the flies were transferred into a second vial and allowed to lay eggs for three more days, before being removed from the vial. Four days after each set of crosses was started, the first set was treated with hydroxyurea and the second set was treated with water. Between the 10^th^ and 18^th^ days, eclosed flies in each vial were scored as homozygous or heterozygous. Relative survival was calculated by dividing the ratio of homozygotes that eclosed in vials treated with hydroxyurea by the ratio of homozygotes that eclosed in respective vials treated with water and converting to a percentage ((%homozygous^treated^/%homoygous^water^)x100)). Mean relative survival for each *rif1*^*null*^ mutant was determined based on the results from 5-10 vials per dose. Vials containing fewer than five flies or two standard deviations below the mean were excluded from the analysis.

In ionizing radiation assays, 20-60 virgin female homozygotes (*rif1*^*null*^*/rif1*^*null*^) were crossed with 10-30 males heterozygous for the same *rif1*^*null*^ allele (*rif1*^*null*^*/ CyO*) in a cage at 25°C. Mated females laid eggs on individual grape juice agar plates and a fresh plate was provided every 24 hours. Once hatched larvae reached the third instar stage, they were exposed to Ce-137 radiation in a Gammator 1000 irradiator. Irradiated larvae were transferred to new bottles that contained standard cornmeal food and were allowed to develop into pupae. Eclosed flies were scored as homozygous or heterozygous and relative survival was calculated by comparing to the mean ratio of homozygotes from two untreated plates (using the same formula above). Flies from three separate plates were counted for each dose of IR and mutant.

### Hatching assays

20-30 virgin female homozygotes (*rif1*^*null*^*/rif1*^*null*^) were crossed with 10-15 male homozygotes of the same genotype in a vial for 24 hours at 25°C. Crosses were transferred to cages at 25°C and allowed to lay eggs for 24 hours on a grape juice agar plate. Unhatched and hatched eggs were counted on each plate 48 hours after the plate was removed from the cross. Assays were repeated in triplicate for each mutant and a Kruskal-Wallis ANOVA was used to determine significance (all statistical tests were done in GraphPad Prism 9).

### *P{w*^*a*^*}* repair assay

*P{w*^*a*^*}*, which contains a *white* gene with an inserted copia retrotransposon, was inserted into the essential *scalloped* gene on the *X* chromosome (Adams *et al*. 2003). Excision of the *P{w*^*a*^*}* element leaves a 14 kb DSB gap that is used to track repair outcomes in Drosophila. A series of crosses was performed to obtain male flies which carried the *P{w*^*a*^*}* allele (*X*^*P{wa}*^*/Y*), were homozygous for the *rif1*^*2*^ allele, and carried a source of a *Δ2-3* transposase on the third chromosome (*P{Δ2-3}99B*, from Bloomington stock #2535). Individual male flies were crossed with four to seven virgin females homozygous for the *P{w*^*a*^*}* allele (*X*^*P{wa}*^*/X*^*P{wa}*^). Repair events were scored in female progeny that carried an intact *P{w*^*a*^*}* allele (*X*^*P{wa}*^*\*/X*^*P{wa}*^) and did not inherit the transposase.

If SDSA occurs at the break, followed by annealing of long terminal repeats, female progeny will inherit a P-element without the copia retrotransposon and will express a dominant *white*^*+*^ allele coding for red eyes. If end-joining occurs, the transposon is lost and female progeny will inherit a paternal *white*^*–*^ allele; the haploinsufficiency of the maternal *P{w*^*a*^*}* element results in yellow eyes. Progeny with apricot eyes were also counted, but this outcome did not distinguish between events in which the *P{w*^*a*^*}* was not excised or precise intersister HR. Welch’s t-tests were used for statistical comparisons.

### *P{EGFP*} repair assay

A plasmid that contains *EGFP* and a Cas9 nickase gene (not relevant to this study) was inserted into the *P{CaryP}attP18* element within *6C12* on the *X* chromosome (*y*^*1*^, *w*^*67c23*^, *P{pBID-3xP3-EGFP-vasa-HACas9*^*D10A*^*-attB}*, gift from Avital Rodal). Initially, crosses were performed at 25°C to obtain *white*^*–*^ males which carried the *P{EGFP}* element (*X*^*P{EGFP}*^*/Y*), were homozygous for the *rif1*^*2*^ allele, and carried the previously mentioned *P{Δ2-3}99B* transposase to excise the *P{EGFP}* element and create a 19 kb gap in the *X* chromosome. Due to low survival of males at 25°C, the crosses were repeated at 18°C. Single males were then crossed with four to seven *w*^*1118*^ virgin females at 18°C and germline repair events were scored in the female progeny.

No excision or full HR using the sister chromatid results in an intact GFP allele and the female progeny have fluorescent-green eyes. End-joining or SSA does not restore the GFP allele and female progeny have white eyes. GFP+ and white-eyed flies were scored for each genotype and statistical comparisons were made with Welch’s t-tests.

### DR-*white.mu* repair assay

The DR-*white*.*mu* reporter distinguishes between pathways utilized to repair I-*Sce*I-induced breaks in Drosophila and was described previously (Do *et al*. 2014). A stepwise cross scheme was used to create males homozygous for the *rif1*^*1-7*^ allele, carrying one copy of the DR-*white*.*mu* reporter, and one copy of an I-*Sce*I endonuclease expression construct driven by a ubiquitin promoter (*P{Ubiq::I-SceI, mw*^*+*^*}*, gift from William Engels). Individual males were crossed with three *yw* females to recover repair events in the next generation.

Male and female progeny from each cross were categorized into three different types of repair events based on their eye and body colors: a y+w+ phenotype indicated repair by intrachromosomal HR, a y+w-phenotype indicated either no cut occurred, end-joining repair, or intersister HR, and a y-w-phenotype indicated repair by SSA. A cross was excluded from an analysis if the total progeny were fewer than two standard deviations below the mean for total progeny among all crosses for a particular genotype. The percentage of progeny in each repair category was plotted for each cross and statistical comparisons were made with Welch’s t-tests.

### Sequencing DR-*white.mu* repair junctions

For analysis of repair junctions from the DR-*white*.*mu* assay, a single male was taken from the progeny of each cross. To extract genomic DNA, each fly was squished in 50 μL of squishing buffer (10 mM Tris-Cl pH 8.2, 1 mM EDTA, 25 mM NaCl, 200 μg/mL of Proteinase K) and incubated at 37°C for 30 minutes, then 95°C for 2 minutes (Gloor *et al*. 1993). The repair junction was amplified using previously validated *Sce*.*white* primers (DR-*white* 1.3 forward, 5’ GTTTTGGGTGGGTAAGCAGG 3’; DR-*white* 1a reverse, 5’ AGACCCACGTAGTCCAGC 3’) (Do *et al*. 2014). PCR products were isolated by gel extraction (Macherey-Nagel) and sequenced with a primer located upstream of the I-*Sce*I cut site (DR-*white*2, 5’ ATGCAGGCCAGGTGCGCCTATG 3’) (Eton Bioscience).

## Results

### Drosophila tolerate replication stress and DNA damage without Rif1

In mammals, RIF1 helps prevent DNA damage by regulating stalled replication forks. Depletion of RIF1 in human cells causes sensitivity to stalled replication intermediates induced by hydroxyurea (Garzón *et al*. 2019) and mouse RIF1 is recruited to stalled forks to prevent fork degradation (Mukherjee *et al*. 2019).

Since Drosophila Rif1 also localizes to replication forks (Munden *et al*. 2018), we tested if Drosophila Rif1 is required to tolerate replication stress induced by hydroxyurea (HU). HU stalls replication by depleting free nucleotide (nt) pools in cells and consequent collapsed replication forks are often converted to DSBs (Krakoff *et al*. 1968; Petermann *et al*. 2010). Interestingly, we found *rif1* mutants were not sensitive to high concentrations of HU (Figure 2A). Overall, these data show Drosophila Rif1 functions differently than mammalian RIF1 and is not required to survive treatment with HU.

**Figure 2.**
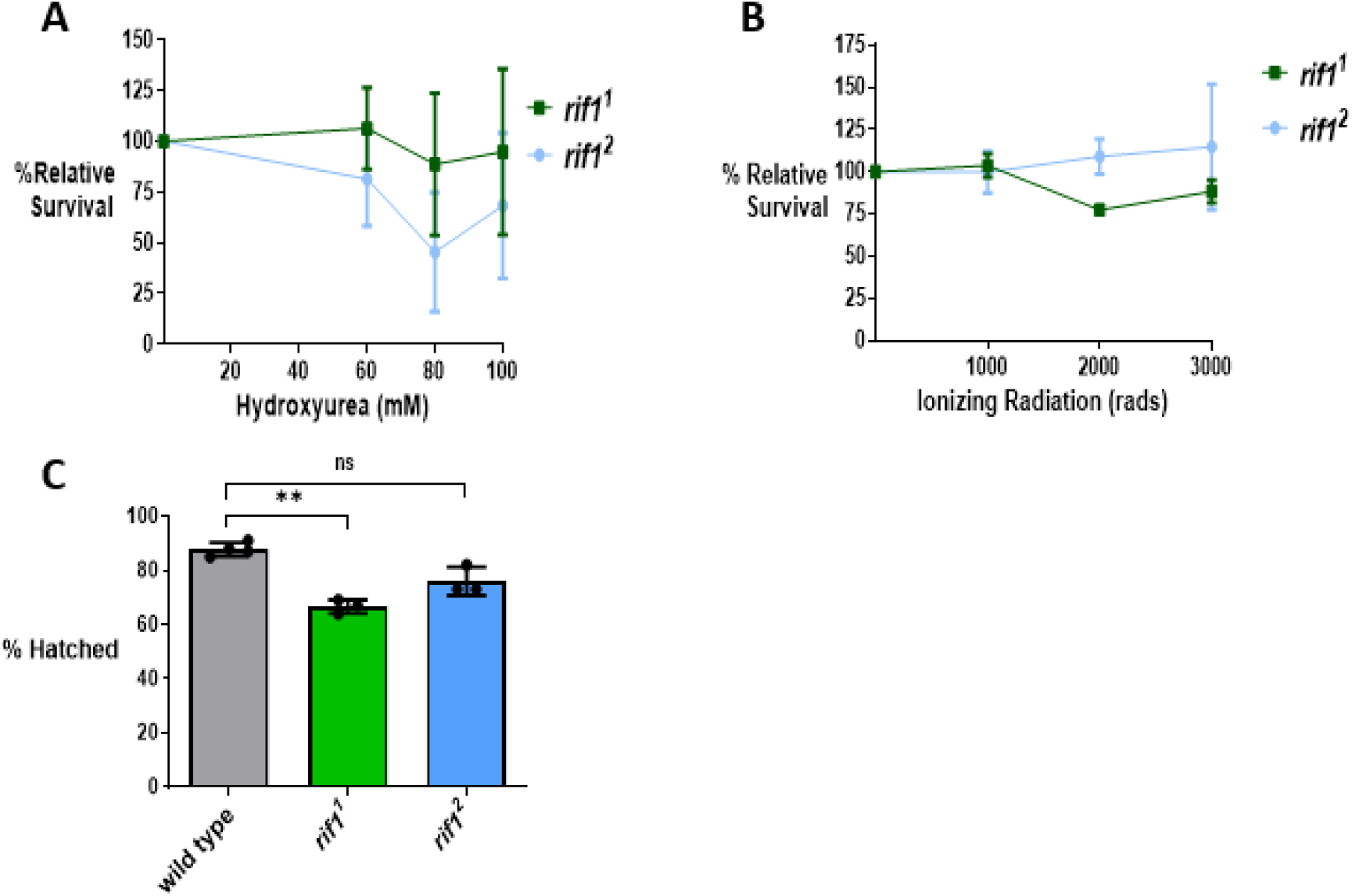
Drosophila Rif1 is not required to resolve replication stress or DNA damage. A) Survival to adulthood of *rif1* mutants treated with hydroxyurea. Relative survival was calculated by dividing the ratio of homozygous flies that eclosed after treatment by the ratio of homozygous flies that eclosed without treatment. Shown are the mean survival and standard deviation for 5-10 replicate crosses per dose. B) Survival to adulthood of *rif1* mutants treated with ionizing radiation relative to untreated larvae. Shown are the mean survival and standard deviation for 3 replicate crosses per dose. C) Percent of *rif1* null homozygous eggs that hatched 48 hours after egg laying. Shown are the mean and standard deviation for 3 replicate counts for each genotype. Hatching percentages were compared via Kruskal-Wallis ANOVA, **p=0.001 to 0.01, ns=not significant.

Multiple lines of evidence also show vertebrate RIF1 localizes to breaks in DNA and is important for cellular survival after exposure to ionizing radiation (IR) (Silverman *et al*. 2004; Chapman *et al*. 2013; Escribano-Díaz *et al*. 2013; Feng *et al*. 2013). RIF1 supports survival by promoting NHEJ at DSBs (Chapman *et al*. 2013; Escribano-Díaz *et al*. 2013) and suppresses γ-H2AX foci from accumulating in cells (Buonomo *et al*. 2009; Chapman *et al*. 2013; Bakr *et al*. 2016; Eke *et al*. 2020). To determine if Rif1 is required to resolve DSBs in Drosophila, we treated *rif1*^*1*^ and *rif1*^*2*^ mutant larvae with increasing doses of ionizing radiation (IR). Surprisingly, neither mutant showed increased sensitivity to IR (Figure 2B).

Although Rif1 is not required for tolerance of exogenous damage, we wondered whether it may play a role in the repair of endogenously-derived DSBs that occur in follicle cells of the developing egg. Within the follicular epithelium, four loci called *Drosophila amplicons in follicle cells* (DAFCs) undergo repeated amplification (re-replication) (Mahowald *et al*. 1979; Claycomb and Orr-Weaver 2005). Re-replication of these regions allows for increased protein production and formation of the eggshell (Spradling and Mahowald 1980), but this rapid replicative process causes numerous DSBs, which are repaired by a variety of processes (Davidson *et al*. 2006; Alexander *et al*. 2015). Previous work in the lab has shown that TMEJ-deficient flies lacking POLQ have thin eggshells and severe hatching defects (Alexander *et al*. 2016). Interestingly, it was shown that *rif1*-mutant eggs exhibit modest and substantial egg hatching defects (Munden *et al*. 2018; Seller and O’Farrell 2018). Since these observed defects may suggest Rif1 has a partial role in repairing breaks that arise during re-replication, we further explored this phenotype by measuring the hatching rates of *rif1* mutants ourselves. *rif1*^*1*^ mutant eggs exhibited significantly lower hatching frequencies than wild type, but the overall decrease in hatching was minor and *rif1*^*2*^ mutants did not exhibit a significant decrease (Figure 2C). A separate *rif1 null* mutant also did not exhibit significant hatching defects and this suggests the decreased hatching observed in *rif1*^*1*^ was independent of Rif1 function (Figure S1B). Additionally, *rif1*-mutant eggshells had normal hexagonal patterning (Turner and Mahowald 1976), without the patchiness observed in *polq* mutants (Alexander *et al*. 2016). Overall, our results suggest Rif1 is not required for the development of the eggshell or to repair DSBs that arise during re-replication in follicle cells.

### Rif1 has a critical role in the repair of P-element-induced DSBs

Since Rif1 regulates DSB repair pathway choice in yeast and mammals by regulating resection, we wondered whether Rif1 may contribute to repair pathway choice in Drosophila. To test this, we used a site-specific gap repair assay called *P{w*^*a*^*}* that distinguishes between different models of repair at P-element-induced breaks (Adams *et al*. 2003). In this assay, a transposase catalyzes excision of the *P{w*^*a*^*}* element from the *X* chromosome and leaves a two-ended DSB with 17 nt-overhangs in the pre-meiotic germline of a parental male (Figure 3A). Break repair by HR can be distinguished from end-joining based on the eye color of female progeny.

**Figure 3.**
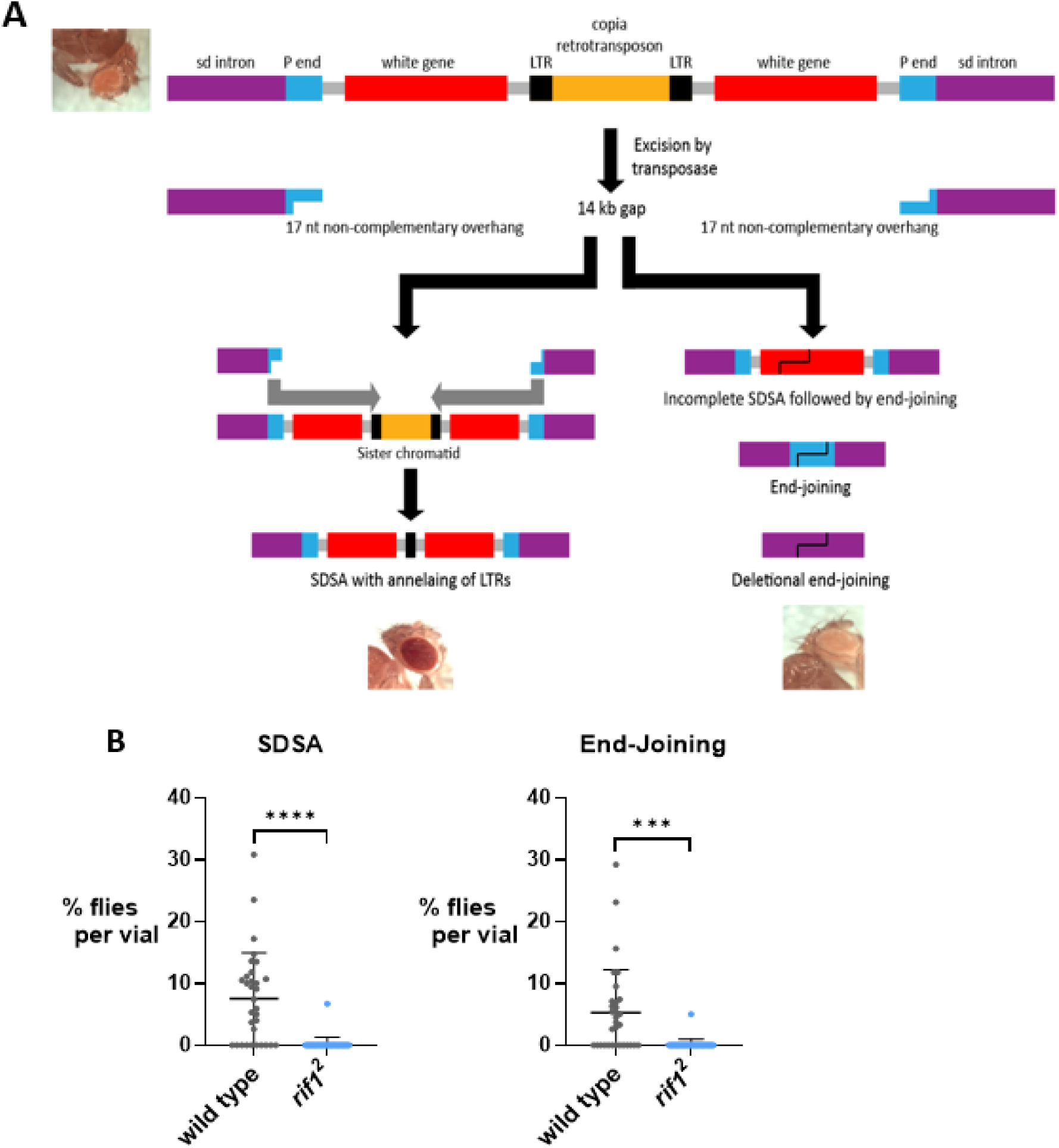
Rif1 is required for the repair of P-element-induced DSBs. A) Diagram outlining the *P{w*^*a*^} assay from Adams *et al*. 2003. The *P{w*^*a*^} construct contains the white gene disrupted by a copia retrotransposon that is flanked by two long terminal repeats (LTR). It is inserted within an intron of the *scalloped* (sd) gene on the *X* chromosome. A P-element transposase excises the *P{w*^*a*^} element to yield a 14 kb gap and two 17 nt non-complementary 3’ overhangs in the male pre-meiotic germline. Following a cross to homozygous *P{w*^*a*^} females, repair events from the male germline are recovered in female progeny *in trans* to an intact copy of *P{w*^*a*^}. Repair by SDSA and annealing at the LTRs produces an uninterrupted *white* gene and yields red-eyed progeny, while repair by end joining or interrupted SDSA followed by end joining yields yellow-eyed progeny. No excision or error-free repair yields apricot-eyed progeny (not shown). B) Homozygous *rif1*^*2*^ mutant males possessing both *P{w*^*a*^*}* and the transposase were crossed with homozygous *P{w*^*a*^} females and eye color of the female progeny was scored. Each data point represents the eye color percentage for progeny from one cross (wild type=36 total crosses, *rif1*^*2*^=32 total crosses). Error bars signify standard deviation in both graphs. ***p=0.0001 to 0.001, ****p<0.0001 (Welch’s t-test).

The *P{w*^*a*^*}* assay was conducted in both wild-type and *rif1*-mutant backgrounds, followed by comparative analysis of the repair pathways utilized in each system (Figure 3B). Strikingly, in the absence of Rif1, repair by HR or end-joining rarely occurred. Further, red patches in the eyes of wild-type parental males indicated somatic repair by SDSA, whereas *rif1*-mutant males rarely exhibited the same mosaicism (Figure S2). Collectively, this suggests that Rif1 is important for the repair of P-element-induced DSBs in the pre-meiotic germline and in the soma.

To validate the repair defects observed in the *P{w*^*a*^*}* assay, we designed and conducted another transposase-induced DSB assay using a P-element transposon marked by GFP (*P{EGFP}*) on the *X* chromosome. Here, a transposon is also excised in the male pre-meiotic germline and the repair mechanism is determined by the eye color of female progeny, following a cross with white-eyed females. End-joining is quantified by the recovery of GFP– female progeny, while HR and no-cutting events cannot be differentiated as they are both represented by GFP+ female progeny (Figure 4A).

**Figure 4.**
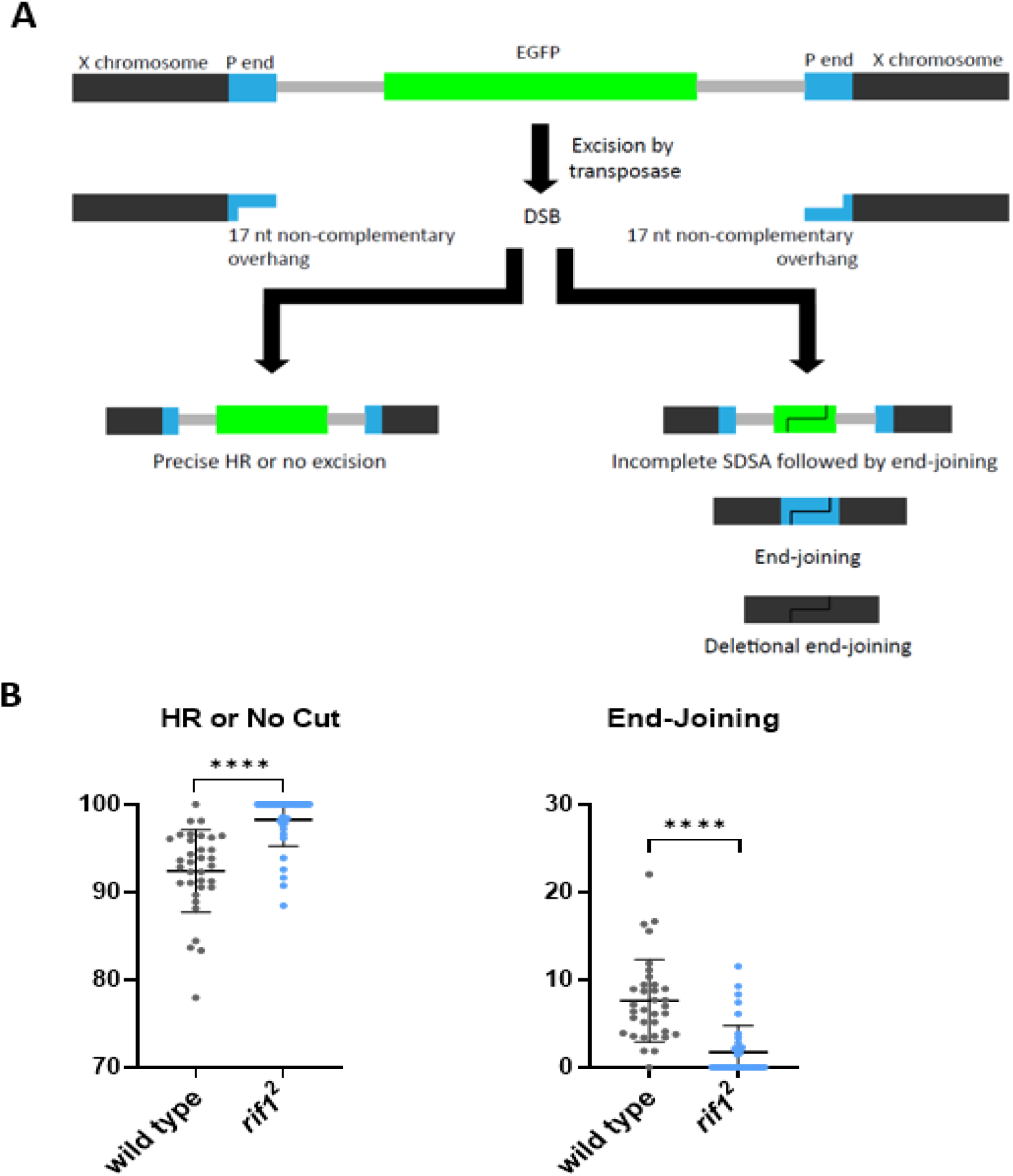
Rif1 is important for repair of P-element-induced DSBs at an additional genomic locus on the *X* chromosome. A) Diagram outlining the *P{EGFP*} assay. Excision of the *P{EGFP}* element by a transposase creates a 19 kb gap with two 17 nt non-complementary 3’ overhangs in the *X* chromosome. Individual male flies hemizygous for *P{EGFP*} and heterozygous for the transposase were crossed with homozygous *w*^*1118*^ females and female progeny eye color was scored. No cut or precise HR yields a full *EGFP* allele and the progeny have fluorescent green eyes. End-joining does not restore the *EGFP* allele and the progeny have white eyes. B) Percent of progeny with no cut or HR repair events (GFP^+^ eyes) and end-joining repair events (white eyes). Each data point represents the eye color percentage for progeny from one cross (wild type=34 total crosses, *rif1*^*2*^=37 total crosses). Eye color percentages for *rif1* mutants were compared to wildtype via Welch’s t-test, ****p<0.0001. Error bars signify standard deviation in both graphs.

We initially conducted this assay at 25°C, but despite extensive screening only five *rif1*-mutant males possessing both the transposon and transposase survived, and only two of those flies were fertile. We surmised that the *P{EGFP}* transposon might excise more frequently than *P{w*^*a*^*}*, leading to sterility and lethality in the absence of repair in *rif1* mutants. We therefore repeated the assay at 18°C to reduce transposase activity. Interestingly, we still observed a significant decrease in the frequency of end-joining in *rif1* mutants relative to wild type (Figure 4B). An increase in fluorescent-green eyes was also observed in progeny derived from the *rif1*-mutant background, but since this phenotype does not distinguish between no cut or HR, we cannot conclude there was an increase in HR in the *rif1* mutants. Like the mosaicism detected in the *P{w*^*a*^*}* assay, non-GFP^+^ patches were detected in the eyes of wild-type parental male flies, but these patches were hardly detectable in the *rif1*-mutant male flies. Together, these results provide strong evidence that Rif1 is needed for the repair of P-element-induced breaks in both the germline and the soma.

### Rif1 is involved in repair pathway choice at I-*Sce*I-induced breaks

We observed Drosophila Rif1 has a critical role in the repair of P-element-induced DSBs, but it is not required to repair IR-induced breaks or breaks that occur in the follicular epithelium of eggs. Therefore, we wondered whether Rif1 functions exclusively at P-element excision sites or if it may also be involved in repair pathway choice at breaks with other types of end structures.

To test this, we induced DSBs using the I-*Sce*I endonuclease, which creates short 4-nt overhangs instead of long 17-nt overhangs produced by P-element excision. We used a previously developed reporter construct called DR-*white*.*mu* (Do *et al*. 2014). The reporter contains an I-*Sce*I recognition sequence and two nonfunctional white genes (*Sce*.*white* and *iwhite)* that flank a yellow transgene (*y*^*+*^) (Figure 5A). Following cleavage by I-*Sce*I, germline repair events are scored in the next generation based on the eye and body colors of the progeny. Repair by intrachromosomal HR restores the missing section of the nonfunctional *Sce*.*white* gene at the cut site and results in a y+w+ phenotype in the progeny. End-joining, intersister HR, or the failure of I-*Sce*I to cut does not restore *Sce*.*white* to a functional white gene and produces a y+w-phenotype. SSA or deletional EJ mechanisms, which require extensive resection, lead to the annealing of *Sce*.*white* and *iwhite* sequences, loss of the yellow transgene, and a y-w-phenotype. Because it was previously shown that the majority of *y-w-* flies contain repair products produced by SSA and not end-joining, we refer to this group as SSA. (Do *et al*. 2014).

**Figure 5.**
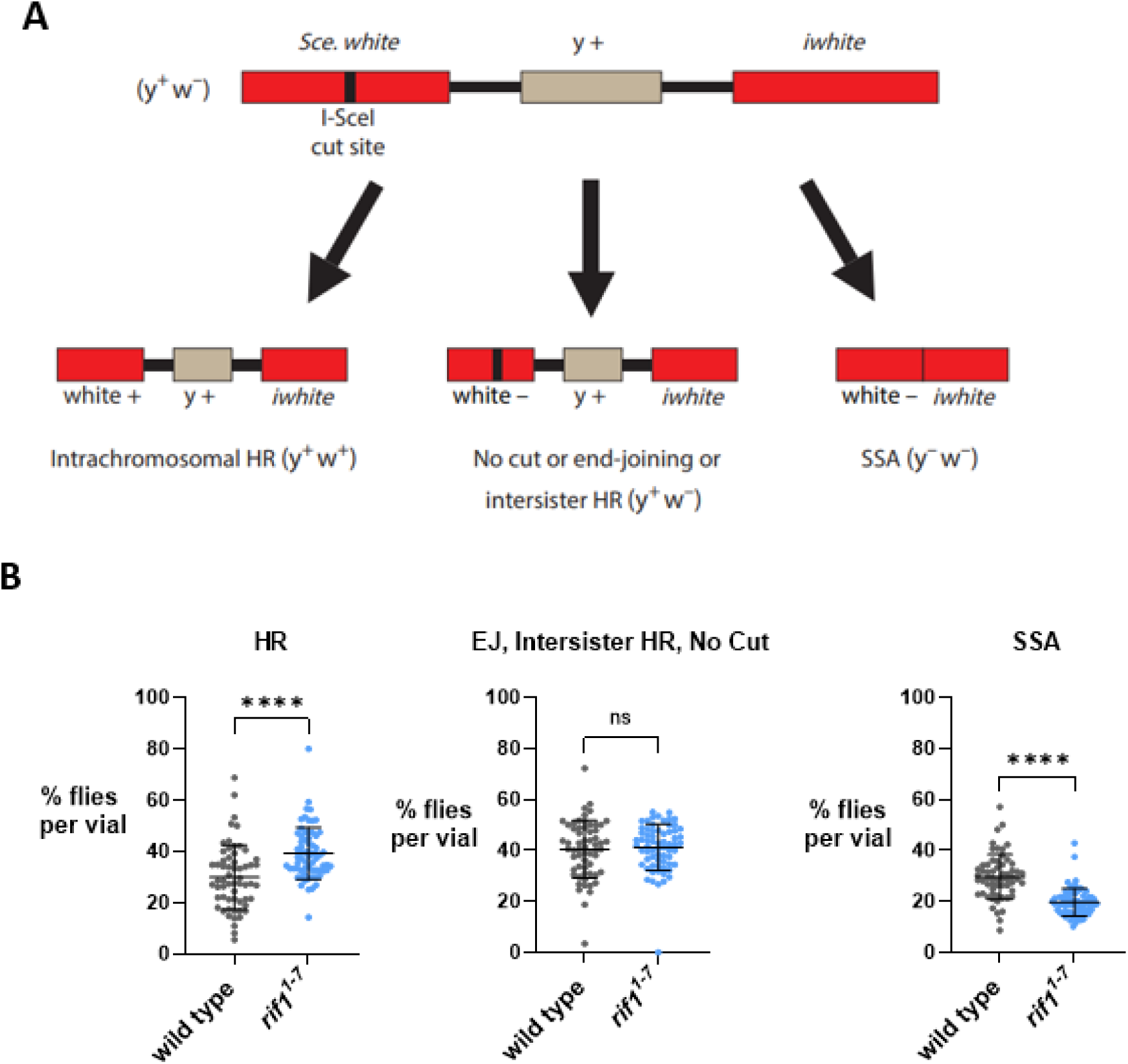
Compensatory shifts in the frequency of HR and SSA in *rif1* mutants. A) Diagram outlining the DR-*white* assay (Do *et al*. 2014). The DR-*white*.*mu* construct contains a white gene disrupted by an I-*Sce*I recognition sequence (*Sce*.*white*), a yellow transgene (y+), and an additional white gene disrupted by 5’ and 3’ truncations (*iwhite*). In addition to the DR-*white*.*mu* sequence, male flies carry a ubiquitously expressed I-*Sce*I construct and are homozygous for either wild-type or mutant (*rif1*^*1-7*^) *Rif1* alleles. Single males are crossed to *yw* females and germline repair events are recovered in the progeny. B) Progeny from wild-type or *rif1*^*1-7*^ single-male crosses were scored for specific repair phenotypes. Each data point represents the occurrence of a repair event in one population of progeny (from one cross; wild type=60 total crosses, *rif1*^*1-7*^=67 total crosses). ****p<0.0001, ns=not significant (Welch’s t-test). Error bars signify standard deviation.

For this assay, we used the *rif1*^*1-7*^ allele, which we recovered from a P-element excision that shifted *Rif1* out-of-frame and removed portions of the 5’ UTR and coding region of the gene (Figure S1A). The *rif1*^*1-7*^ stock behaves similarly to other *rif1* null stocks since the hatching rate is not significantly different from the rates of *rif1*^*1*^ and *rif1*^*2*^-mutant eggs (Figure S1B). After a series of crosses, we obtained healthy *rif1*^*1-7*^ male flies expressing the I-*Sce*I endonuclease. This contrasts with the lethality we observed in transposase-expressing *rif1*-mutant males in the *P{EGFP}* assay. Upon collecting repair phenotypes in the progeny, we observed significantly more HR occurred in *rif1*^*1-7*^ mutants than wild type (Figure 5B). In contrast, SSA occurred less often in *rif1*^*1-7*^ mutants. We did not observe a significant difference between the number of y+w-progeny found in *rif1*^*1-7*^ and wild-type crosses. However, the y+w-phenotype fails to distinguish between the occurrence of end-joining, intersister HR, or no cut. Accordingly, we took one y+w-fly from each cross and sequenced individual repair junctions. Most sequences exhibited cutting by I-*Sce*I and interestingly, both wild-type and *rif1*^*1-7*^-repair junctions contained small 1-10 bp deletions or insertions (Table 1). Thus, with or without Rif1, error-prone end-joining mechanisms often mediate repair of I-*Sce*I-induced breaks. Taken together, our results suggest Rif1 plays a significant role in the choice between repair pathways at I-*Sce*I-induced breaks, by suppressing HR or facilitating deletional repair by SSA.

**Table 1:**
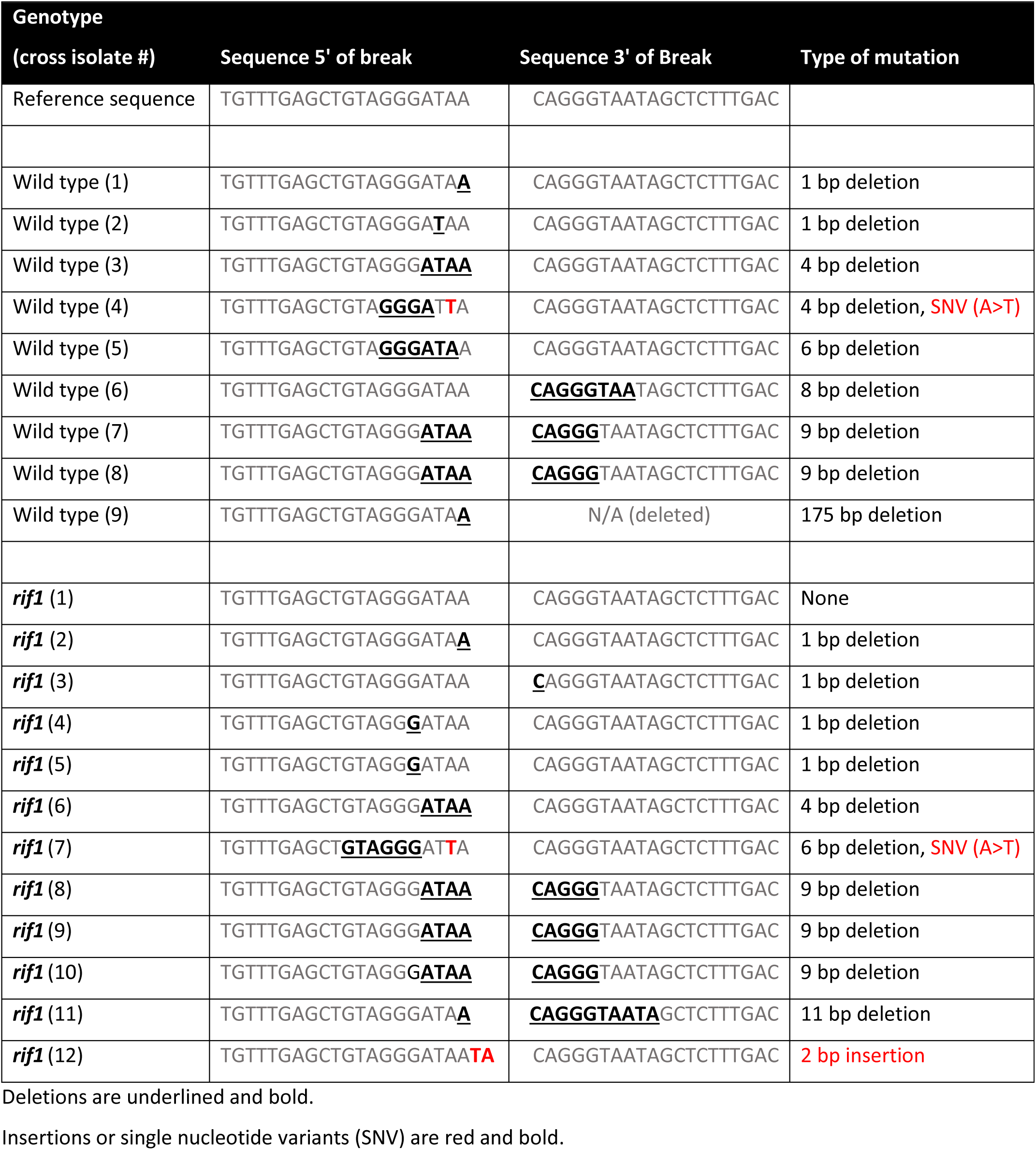
Sequences of wild-type and *rif1*-mutant DR-*white*.*mu* repair junctions.

## Discussion

DSB repair pathway choice begins with the decision to initiate 5’ to 3’ resection. Studies have identified Rif1 as a determinant of DSB repair pathway choice in yeast and mammals via the regulation of resection. Interestingly, in mammals, RIF1 promotes NHEJ by suppressing resection required for HR, while in yeast, Rif1 can both promote NHEJ and facilitate long-range resection to promote SSA. Given these diverse functions of Rif1, we explored the roles of Drosophila Rif1 in repair pathway choice to provide new insights for how Rif1 regulates repair across eukaryotes. Our findings show that although Drosophila Rif1 is not required to repair all DSBs, Rif1 is critical for repair at P-element-induced DSB ends. Additionally, Rif1 suppresses HR at I-*Sce*I-induced breaks but promotes SSA. Collectively, these results suggest that like mammalian RIF1, Drosophila Rif1 partially suppresses HR and like yeast Rif1, Drosophila Rif1 promotes SSA at extensively resected DSB ends. Unique to this study, our genetic data also suggest an essential role for Rif1 in the repair of P-element-induced DSBs.

### Rif1 is not required to resolve replication stress or DNA damage for Drosophila survival

Surprisingly, the absence of Rif1 did not affect the overall survival of Drosophila upon exposure to hydroxyurea or ionizing radiation. These findings differ from several reports in which vertebrate Rif1 promotes survival by resolving replication stress and DNA breaks induced by IR (Silverman *et al*. 2004; Buonomo *et al*. 2009; Chapman *et al*. 2013; Escribano-Díaz *et al*. 2013; Garzón *et al*. 2019; Mukherjee *et al*. 2019). In contrast, these results are consistent with findings in yeast, where the absence of Rif1 does not cause sensitivity to radiomimetic-induced damage or MMS (Martina *et al*. 2014).

Previously, it was shown Drosophila *rad51 (spn-A)* mutants are sensitive to IR, but *lig4* mutants are not sensitive (Mcvey *et al*. 2004). This suggests HR, and not classical NHEJ, is the predominant method for repair at IR-induced breaks in Drosophila. On the other hand, NHEJ is the preferred method for repair of IR-induced breaks in mammals (Ackerson *et al*. 2021). This difference between Drosophila and mammalian repair preferences may partially explain the lack of sensitivity we observed in *rif1* mutants to DNA damaging agents.

Further, Rif1 was not required for egg hatching and *rif1-*mutant females produced eggs with normal eggshells, which suggests it is not essential to repair damage that arises during re-replication in follicle cells. Re-replication occurs at loci important for eggshell assembly, stalls fork progression, and produces DNA damage (Alexander *et al*. 2015). Lig4 helps facilitate fork progression during re-replication, but is only required for preventing eggshell defects and promoting egg hatching when TMEJ is disrupted (Alexander *et al*. 2015; Alexander *et al*. 2016). Two other studies reported 55% and 88% hatching rates for Drosophila *rif1* mutants (Munden *et al*. 2018; Seller and O’Farrell 2018). We observed 66%, 76%, and 85% hatching rates for *rif1*^*1*^, *rif1*^*2*^, and *rif1*^*1-7*^ mutants, respectively (Figures 2C, S1B). Since only one rate significantly differed from wild type, we attribute these slight differences to natural genomic variation among the stocks. Thus, in addition to Lig4 being dispensable for repair during re-replication, Rif1 is not required to repair damage in follicle cells.

### Rif1 is critical for repair of P-element-induced breaks

Given Rif1 helps both yeast and mammals survive DNA damage, our initial findings that Rif1 is not required for survival after DNA damage in Drosophila was surprising. However, because Rif1 has also been shown to impact repair pathway choice in other organisms, we conducted site-specific repair assays to examine this possibility.

Strikingly, in the *P{w*^*a*^*}* assay, we observed both HR and end-joining at P-element-induced breaks are dependent on Rif1. To validate this finding, we excised a different P-element (*P{EGFP}*) on the *X* chromosome and initially observed male infertility. We anticipated the infertility may be due to more excision of *P{EGFP}* relative to *P{w*^*a*^*}* at 25°C, causing an overabundance of DSBs due to failed repair within *rif1*-mutant tissues. Upon lowering the temperature to 18°C, we recovered fertile males and validated the large decrease in end-joining observed in the *P{w*^*a*^*}* assay. These results suggest that a defect in P-element excision was likely not responsible for the decrease in repair. Instead, the most likely interpretation is that repair is severely impaired at P-element-induced breaks in *rif1* mutants.

Site-specific P-element repair assays are a unique assay system to study pathway choice because they produce 17-nt non-complementary overhangs at DSBs. These overhangs mimic short tracts of resected DNA at natural DSBs caused by exogenous or endogenous sources of damage. Since *rif1* mutants did not exhibit severe repair deficiencies in the other assays we conducted, our results suggest that Rif1 is needed to repair breaks with overhangs that are poor substrates for end-joining.

Recently, it was shown that specific residues within the N-terminus of yeast Rif1 interact with DNA and the N-terminus preferentially binds DNA substrates with 30-nt overhangs (Mattarocci *et al*. 2017). This interaction stabilizes the DNA and is also important for the ability of Rif1 to inhibit resection and promote NHEJ. In agreement with this model, the inability of Drosophila *rif1* mutants to repair transposase-induced breaks also suggests Drosophila Rif1 stabilizes ssDNA for repair.

Drosophila Rif1 may protect long overhangs through several different mechanisms. First, Rif1 may prevent machinery involved in long-range resection from binding to short-resected tracts. In yeast, Rif1 prevents binding of two primary players in resection, Mre11 and Dna2, to HO-induced breaks (Martina *et al*. 2014). Mammalian Rif1 prevents RAD51, BRCA1, and CtIP from accumulating at damage (Feng *et al*. 2013; Isobe *et al*. 2021). In our assays, the 17-nt overhangs at P-element-induced breaks are ideal substrates to initiate and increase the efficiency of long-range resection mechanisms (Cejka 2015). Overactivation of long-range resection commonly leads to extensive genomic instability (Tomimatsu *et al*. 2017) and may be responsible for the abrogated repair we observe in Drosophila *rif1* mutants that experience transposase-induced damage. Second, Rif1 may bind the overhangs to stabilize them for repair by preventing degradation. The N-terminus of yeast Rif1 forms dimers and wraps around DNA (Mattarocci *et al*. 2017). This interaction prevents the destabilization of short overhangs at an HO-induced cut. Interestingly, this N-terminal region can be identified in the Drosophila Rif1 protein (Figure S3). In addition, mammalian Rif1 protects stalled replication forks from degradation by DNA2-WRN (Garzón *et al*. 2019). Drosophila Rif1 may work in a similar fashion to both Rif1 orthologs and be especially important to protect long P-element-induced overhangs from degradation.

### Rif1 has subtle repair functions at I-*SceI*-induced breaks, but is important for SSA following extensive resection

The P-element excision assays showed a reliance on Rif1 for repair of breaks with 17-nt overhangs. However, at I-*Sce*I-induced breaks in the DR-*white*.*mu* assay, we observed a significant increase in intrachromosomal HR and a decrease in SSA in *rif1* mutants. The increase in HR in this assay contrasts with the HR defect we observed in the *P{w*^*a*^*}* assay. We posit this difference may be explained by several mechanisms. First, the 4-nt overhangs in the *DR-white*.*mu* assay and the 17-nt overhangs in the *P{w*^*a*^*}* assay may activate different initial mechanisms for repair. In yeast, Rif1 was shown to inhibit checkpoint activation at ssDNA and *rif1* mutants arrest in G2/M upon the accumulation of ssDNA (Xue *et al*. 2011). Since frequent P-element excision also generates ssDNA in the pre-meiotic germline or soma, persistent checkpoint activation may explain why repair failed in *rif1* mutants in the P-element assays. On the other hand, at I-*Sce*I-induced breaks Rif1 may function more like mammalian Rif1 and partially suppress HR (Isobe *et al*. 2021). Second, the increase in HR may be an artifact of the different repair templates used in the two assays. The DR-*white*.*mu* assay detects HR using an intrachromosomal template, while the *P{w*^*a*^*}* assay detects SDSA using an interchromosomal template. Evidence using a modified DR-*white* construct indicates intrachromosomal templates are highly preferred for HR over interchromosomal templates in the pre-meiotic germline (Fernandez *et al*. 2019). Thus, it is possible the intrachromosomal repair template allows for more HR in *rif1* mutants in the DR-*white*.*mu* assay that cannot be seen in the *P{w*^*a*^*}* assay.

Consistent with findings in yeast, we observed a significant decrease in SSA in *rif1* mutants (Martina *et al*. 2014). This suggests Rif1 is either important to perform long-range resection in flies that is necessary for SSA, or it participates in the annealing of direct repeats during SSA. The former was proposed in yeast, but it is unclear if Rif1 is an accessory unit for nucleolytic degradation of DNA or if it protects ssDNA after resection (Martina *et al*. 2014). If it protects ssDNA, it may also assist in the annealing of distal strands for SSA. Currently, we are unable to distinguish between these two possibilities.

In summary, we have uncovered two separate functions for Drosophila Rif1 in this study: a critical role in processing P-element-induced breaks for repair and an additional role in repair pathway choice at I-*Sce*I-induced breaks. We suspect Rif1 may be required for repair of P-element-induced breaks by protecting ssDNA overhangs that are poor substrates for end-joining. This is also the first study to show Rif1 can promote SSA at extensively resected ends in a eukaryote other than yeast and demonstrates Rif1 is not unifunctional at DSB ends. Taken together, our work exhibits Drosophila Rif1 has some unique repair functions relative to other Rif1 orthologs, but also has similar functional tendencies in determining repair pathway choice. Additional mechanistic investigation into the alternative functions we identified in this study will yield new insights into the involvement of Rif1 in DSB repair across eukaryotes.

## Supporting information

Blanch et al supplemental data

## Limitations of the study

Our work supports one or more roles for Drosophila Rif1 in DSB repair. However, our understanding of the mechanisms by which it performs these roles is limited for a few reasons. First, sensitivity assays rely on the failure to repair breaks leading to organismal lethality and as such they are unable to reveal minor roles in repair. If Rif1 is only required to repair a small subset of the heterogeneous breaks induced by IR, a DSB repair role could be missed. Second, because the site-specific repair assays are not always able to assign a specific repair mechanism to each phenotypic outcome, we are somewhat limited in the conclusions that we can make. Third, although our site-specific repair assays are valuable to elucidate *in vivo* differences in repair pathway choice in the presence and absence of Rif1, they do not identify the mechanism by which these differences occur. Understanding the role of Rif1 function at DSBs will require the use of sophisticated biochemical and genomic techniques which can be used to further establish a mechanism.

## Data Availability

Drosophila stocks and mutant gene maps are available upon request. The data required to confirm the conclusions of this study are fully provided within the figures and tables of the article.

## Author Contributions

JRB conceived this study and created the *rif1*^*1-7*^ mutant. JRB, MK, and MM contributed to experimental design. JRB and MK conducted the sensitivity assays. MK conducted the hatching and transposon assays. JTZ, MK, and MM conducted the DR-*white*.*mu* assay. JRB and JTZ prepared the DR-*white*.*mu* repair samples for sequencing. JRB made the figures and conducted the statistical analyses. JRB, MK, and MM wrote the manuscript.

## Acknowledgments

We thank Jan LaRocque, Avital Rodal, William Engels, and Jared Nordman for generous donations of fly stocks. We also thank Jared Nordman for helpful comments on the manuscript and Sara Martin, Tyler Maclay, and other members of the McVey lab for valuable discussions and contributions to the project. This work was funded by grant MCB1716039 from the National Science Foundation and grant R01GM125827 from the National Institutes of Health.

