## Supplementary material for "Drosophila Rif1 is critical for repair following P-element excision and influences pathway choice at double-strand breaks": Blanch et al supplemental data

Drosophila Rif1 engages in break repair and pathway choice

### Supplementary Section

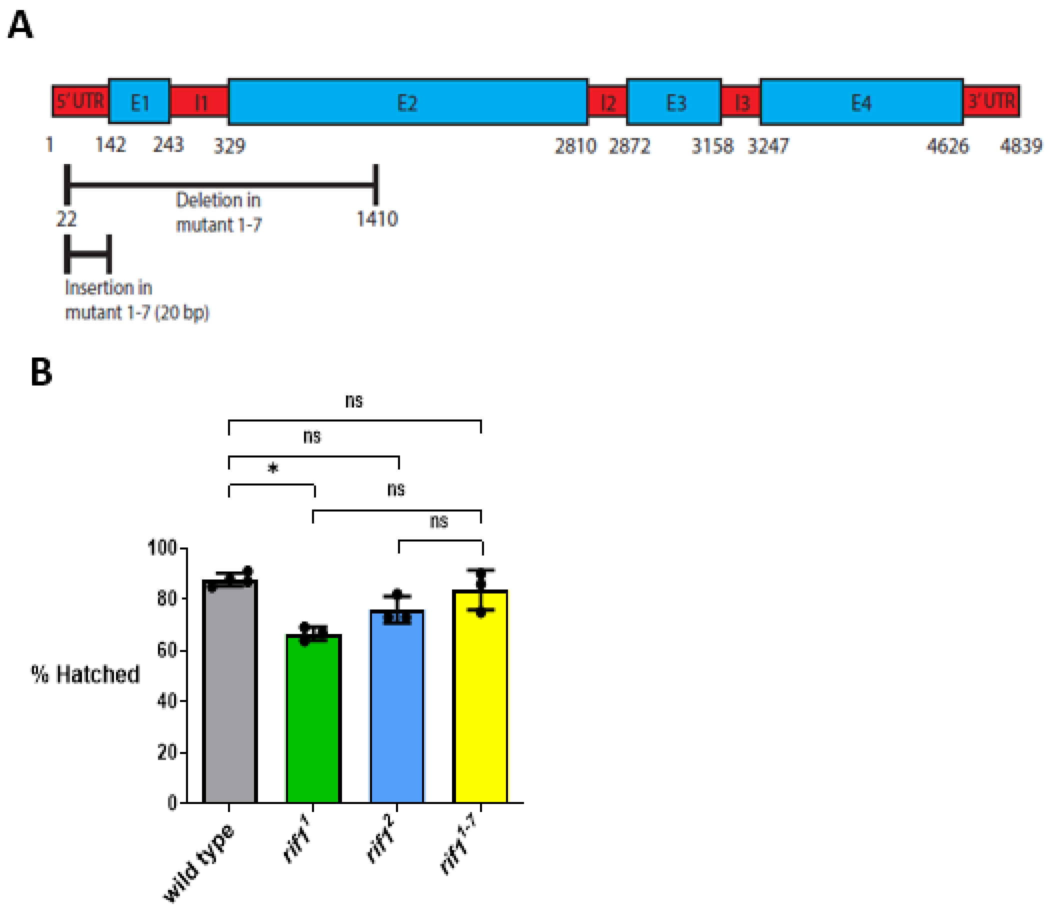


**Figure S1.** A) *Rif1* gene map and locations of mutations in the *rif1*$^{1-7}$ *null* mutant. E=exon, I=intron, UTR=untranslated region. A P-element excision led to a 1,389 bp deletion and 20bp insertion in *Rif1*. The deletion removed 120 bp of the 5' UTR and 1184 bp of the *Rif1* coding region and shifted the entire *Rif1* coding sequence out-of-frame. B) Hatching frequency for *rif1*$^{1-7}$ mutant relative to other *rif1*$^{null}$ mutants. Hatched eggs were counted 48 hours after flies were removed from the cages. Shown are the mean and standard deviation for 3 replicate counts for each genotype. Hatching percentages were compared via Kruskal-Wallis ANOVA,*p=0.01 to 0.05, ns=not significant.

Drosophila Rif1 engages in break repair and pathway choice

Justin R. Blanch*[1$], Manan Krishnamurthy*[1,2], Jacob T. Zuckerman[1], Mitch McVey[1]

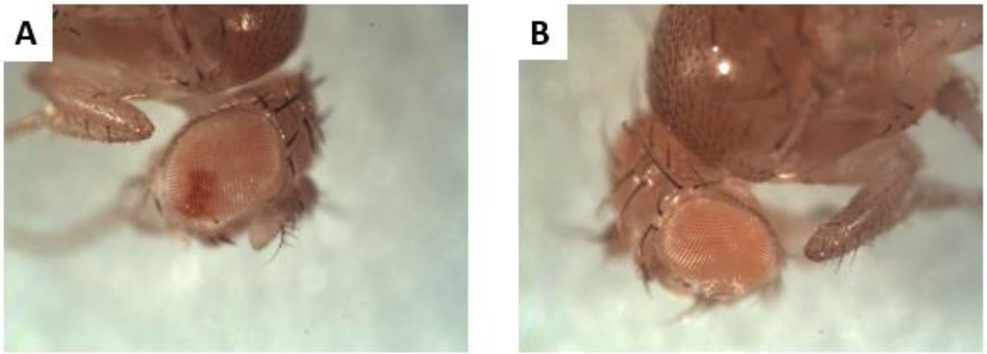


**Figure S2.** Mosaicism detected in parental males carrying the *P{w$^a$}* allele and the transposase in the *P{w$^a$} assay*. A) Red patches that indicated somatic repair by SDSA were often observed in the eyes of wild-type males. B) *rif1*-mutant males rarely exhibited red mosaicism and had apricot-colored eyes (no somatic repair).

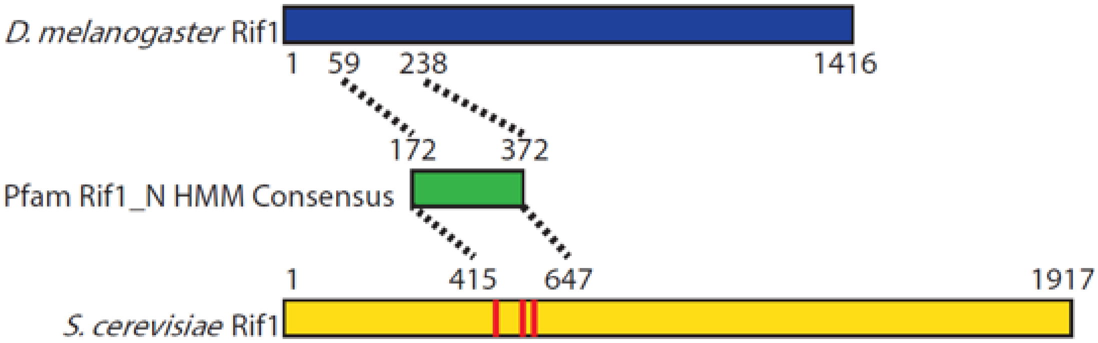


**Figure S3.** Identification of Pfam Rif1_N consensus region in *D. melanogaster and S. cerevisiae* Rif1 protein sequences. The protein sequence for each ortholog were scanned separately for Pfam domain clans (https://pfam.xfam.org/). Regions within the N-termini of both Rif1 orthologs matched the Rif1_N HMM (Hidden Markov model) consensus sequence. These HMM regions were realigned in SnapGene 5.1.7 to identify the Pfam consensus sequence that is found in both proteins (172-372). This shows that the N-termini of both proteins are related to the same functional clan. Red bars in the yeast protein represent three residues (K437, K563, K570) in the consensus region that were previously shown to help stabilize DSB ends for repair (Mattarocci *et al.* 2017).
